# Expectation pooling: An effective and interpretable pooling method for predicting DNA-protein binding

**DOI:** 10.1101/658427

**Authors:** Xiao Luo, Xinming Tu, Yang Ding, Ge Gao, Minghua Deng

**Author notes:** The first two authors should be regarded as Joint First Authors.

## Abstract

**Motivation:** Convolutional neural networks (CNNs) have outperformed conventional methods in modeling the sequence specificity of DNA-protein binding. While previous studies have built a connection between CNNs and probabilistic models, simple models of CNNs cannot achieve sufficient accuracy on this problem. Recently, some methods of neural networks have increased performance using complex neural networks whose results cannot be directly interpreted. However, it is difficult to combine probabilistic models and CNNs effectively to improve DNA-protein binding predictions.

**Results:** In this paper, we present a novel global pooling method: expectation pooling for predicting DNA-protein binding. Our pooling method stems naturally from the EM algorithm, and its benefits can be interpreted both statistically and via deep learning theory. Through experiments, we demonstrate that our pooling method improves the prediction performance DNA-protein binding. Our interpretable pooling method combines probabilistic ideas with global pooling by taking the expectations of inputs without increasing the number of parameters. We also analyze the hyperparameters in our method and propose optional structures to help fit different datasets. We explore how to effectively utilize these novel pooling methods and show that combining statistical methods with deep learning is highly beneficial, which is promising and meaningful for future studies in this field.

**Supplementary information:** All code is public in https://github.com/gao-lab/ePooling

## 1 Introduction

DNA-binding proteins play important roles in gene regulation. The transcription of each gene is controlled by a regulatory region of DNA placed relatively near the start of the transcription site. Several experimental methods, such as ChIP-Seq(Zhang *et al.*, 2008), have been proposed to detect protein-DNA bindings in vivo. Convolutional neural networks (CNNs) have been successfully used to identify functional motifs in massive genomic databases(Alipanahi *et al.*, 2015; Zhou and Troyanskaya, 2015). Analogous to the computer vision task for two label image classification, genomic sequences are first encoded (in one-hot format); then, the 2-D convolution operation is transformed into a 1-D convolution with 4 channels. Following a convolutional layer, pooling layers have been widely applied as effective feature extractors to 1) reduce the feature size and 2) gain invariance to small input transformations to increase model robustness.

While multiple pooling strategies have been proposed(Lu *et al.*, 2015; Lee *et al.*, 2016; Zeiler and Fergus, 2013; Zhai *et al.*, 2017; Huang *et al.*, 2018; Graham, 2014; He *et al.*, 2014; Gulcehre *et al.*, 2014; Xie *et al.*, 2015), max pooling and average pooling are popularly utilized in practical models(LeCun *et al.*, 1990, 1998; Boureau *et al.*, 2008; Jarrett *et al.*, 2009). Max pooling is done by applying a max filter to subregions of the initial representation and global pooling utilizes a average filter. It has been theoretically shown that max pooling improves discriminability over average pooling(Boureau *et al.*, 2010) though max pooling causes overfitting easily. Global max pooling(Lin *et al.*, 2013) are mostly utilized in models of motif detecting, because it has a reasonable statistical meaning of choose the biggest score after convolution. However, max pooling which loses some information may be not optimal and lack of the better interpretation of probability(Alipanahi *et al.*, 2015) on motif inference.

Inspired by the expectation-maximization (EM) algorithm, we propose a new global pooling method: expectation pooling. Evaluations on both simulated and real-world data demonstrate that expectation pooling improves motif identification performance significantly. We further analyze the hyperparameters used in our method and propose optional structures to help fit different datasets. Expectation pooling is both mathematically sound and provides a plausible statistical interpretation for a CNN. All the code used to implement expectation pooling and reproductions of all figures in the manuscript are publicly available at https://github.com/gao-lab/ePooling.

## 2 Materials and methods

### 2.1 Detecting sequence motifs with CNN

Our baseline neural network architecture for motif detection is the simplest model, without a fully connected layer (see the Supplementary Notes), as shown in Figure 1(a). The inputs are DNA sequences; however, the neural network model requires numerical input. Consequently, each sequence is transformed into a one-hot format. Specifically, the sequences are transformed into 4 × *L* matrices where each base pair in a sequence is denoted as one of four one-hot vectors [1, 0, 0, 0], [0, 1, 0, 0], [0, 0, 1, 0] and [0, 0, 0, 1]. The first layer is a 1-D convolutional layer with ReLU activation(Radford *et al.*, 2015), which can be considered as a motif scanner. The second layer is our expectation pooling layer, which will be discussed in the next section. The last layer is a fully connected layer with one output. We use sigmoid activation to obtain the probability of a sample being positive.

**Fig. 1.**
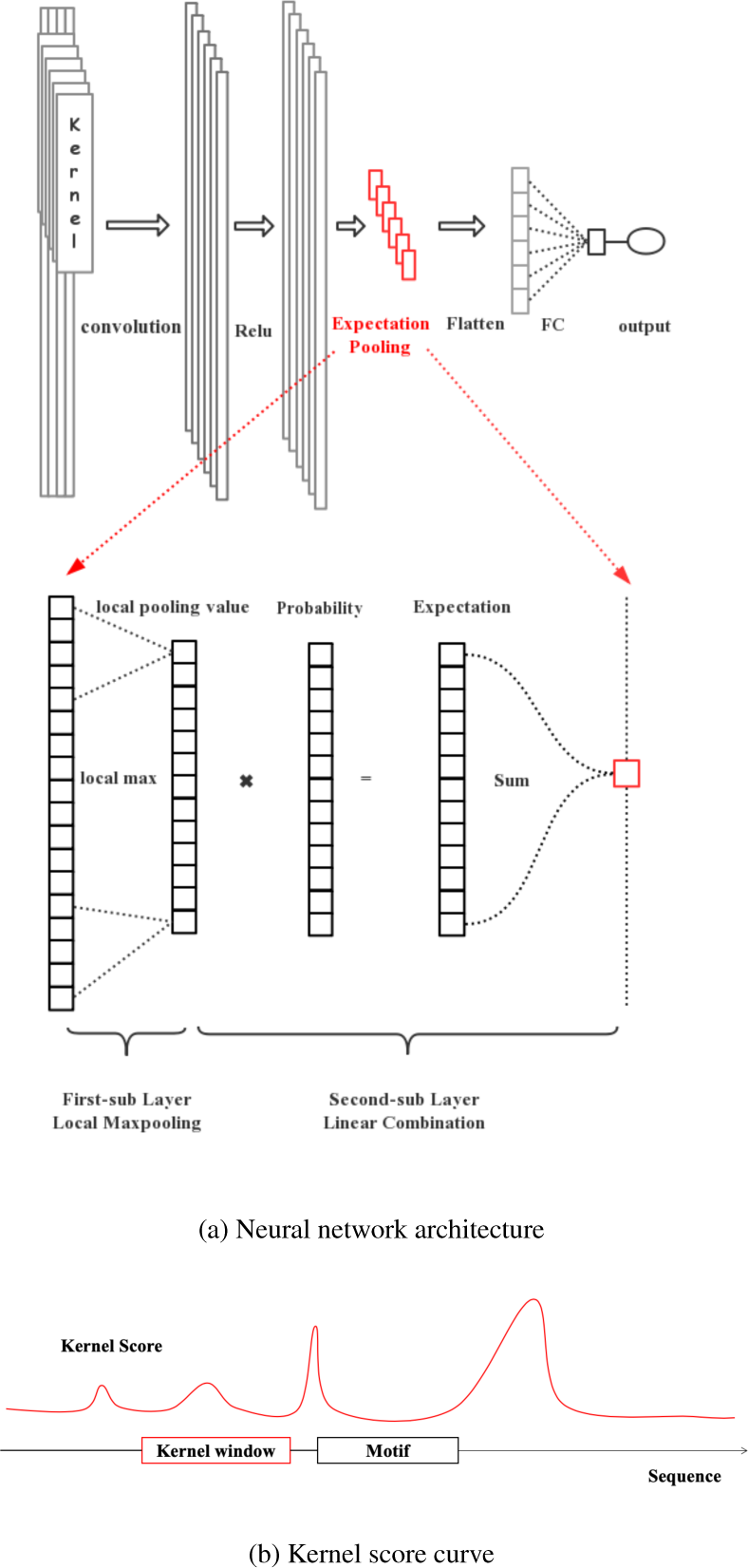
(a): The first layer is a convolutional layer followed by a ReLU activation function. The next layer is the proposed expectation pooling, which is explained in the next section. The third layer is a dense layer that linearly combines the outputs of all the kernels. The last is a sigmoid function that converts the values obtained in the dense layer to a probability between 0 and 1. The expectation pooling layer architecture is as follows: First, we use a local max pooling to filter noise. Then, we calculate the probability at each position and obtain the approximate expectation of the log probability using Equation 3. (b): After calculating the distribution of latent variables, the red curve is the score curve, which is equivalent to the curve of the probability that some positions are motif starting points.

### 2.2 Expectation pooling

Briefly, expectation pooling calculates the weighted average of locally max pooled values. Specifically, the expectation pooling consists of two sublayers: the first sublayer is a 1-D local max pooling with a window size of *q* with zero padding, and the output length is 1*/q* of the origin length. The second sublayer is a dense layer of size 1 with nonparameterized weights, which ensures that the whole pooling layer has no additional parameter. Overall, the output of the expectation pooling layer is a weighted linear combination of the larger part of its input; the tendency is that for larger input values result in larger weight assignments. The mathematical formula is shown below:

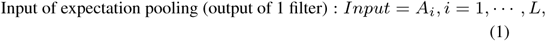

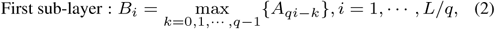

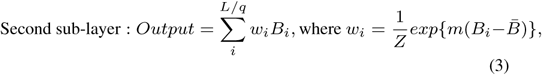

where 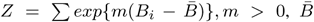 refers refers to the average of all scores of the layer, and *m* and *q* are hyperparameters determined by validation. The *l*_1_ penalty is added to the weights to help the model to assign insignificant input features with zero weight and avoid overfitting(Tibshirani, 1996). If necessary, zero padding can be added to the end of *A*_*i*_.

### 2.3 Implementation of the parameterized convolutional neural networks

The hyperparameters for simulated datasets in the convolutional layer include the number and length of convolution kernels, the number of epochs, the training batch size and the optimizer [see Table 1]. The hyperparameters in the pooling layer include the standard window width for local max pooling and the weight-scale parameter *m* used for expectation pooling (if *m* is not learnable).

**Table 1.** CNN parameter settings for the simulated datasets

| Name | Possible Values |
| --- | --- |
| kernel number | 128 |
| kernel length | 24 |
| training batch size | 32 |
| number of epoch | 100 |
| optimizer | Adam($lr = 0.01, \beta_1 = 0.9, \beta_2 = 0.999, \epsilon = None, \text{decay} = 0.0, \text{amsgrad} = False$) |
| random seed | 1, 2, 3, 4, 5, 6, 7, 10, 15, 20, 30, 40, 50, 100, 200, 300, 400, 500, 600, 1000 |

For training, we used cross-entropy as a loss function without any weight decay, and the models were trained using the standard error backpropagation algorithm and the Adam optimizer(Kingma and Ba, 2014).

We utilized the area under the ROC (AUC)(Davis and Goadrich, 2006; Fawcett, 2004) to assess the prediction performance. Our model is implemented using Keras (Chollet *et al.*, 2015) for Python.

### 2.4 Datasets

#### 2.4.1 Simulated dataset

For the simulations, we used the TRANSFAC(Wingender *et al.*, 1996) database to evaluate whether expectation pooling improves model performance. Each simulated dataset includes both negative and positive samples (i.e., sequences). Each negative sample consists of i.i.d. nucleotides conforming to a multinomial distribution with a probability of 0.25 for each of *{A, C, T, G}*. A positive sample is constructed the same way as a negative sample except that sequences from certain motif(s) are inserted at random locations. Specifically, the sequences inserted in the positive samples for each of the three simulated datasets are as follows:

- **simulated dataset 1**: a sequence generated from the first, shorter motif;
- **simulated dataset 2**: a sequence generated from the second, longer motif;
- **simulated dataset 3**: a sequence generated from either the first or the second motif; the choice of motif for each positive sample is determined randomly with equal probability.

It should be emphasized that **simulated dataset 3** is an important pattern in omics data: a protein is likely to bind to more than one motif in the DNA sequence.

#### 2.4.2 Real dataset

We chose the 690 ChIP-seq ENCODE datasets tested by the DeepBind model(Alipanahi *et al.*, 2015). Each of these datasets corresponds to a specific DNA-binding protein (e.g., transcription factor); its positive samples are 101 bp DNA sequences that were experimentally confirmed to bind to a given protein, and its negative samples were created by shuffling the positive samples. All the datasets were downloaded from http://cnn.csail.mit.edu/.

## 3 Results

### 3.1 Expectation pooling performs better than global max or average pooling on this simulated data

In this section, we compare expectation pooling with global max and average pooling on the simulated datasets. We first selected the simplest CNN model with no hidden layers, with *m* = 1, local window size = 10, batch size = 32, kernel length = 24, and kernel number = 128 (the same specifications listed in Table 1). Immediately after the convolutional layer we appended one of the three different pooling methods above to assess which pooling method achieved a better performance. We set several random seeds to evaluate the robustness of the model’s performance on the simulated datasets.

We found that compared to global max or average pooling, expectation pooling improved the motif finding performance on all three simulated datasets [Figure 2](a)]. Specifically, the model with expectation pooling resulted in a considerable reduction in accuracy variance (measured by AUC) than did the models with global max or average pooling, which suggests that it is more robust to different random seeds. Moreover, we found that the difference between training loss and testing loss was still moderate for the expectation pooling-based model after tens of epochs and it was smaller than that of the models with global max/average pooling, further suggesting that less overfitting occurred during training (see Figure 2](b)).

**Fig. 2.**
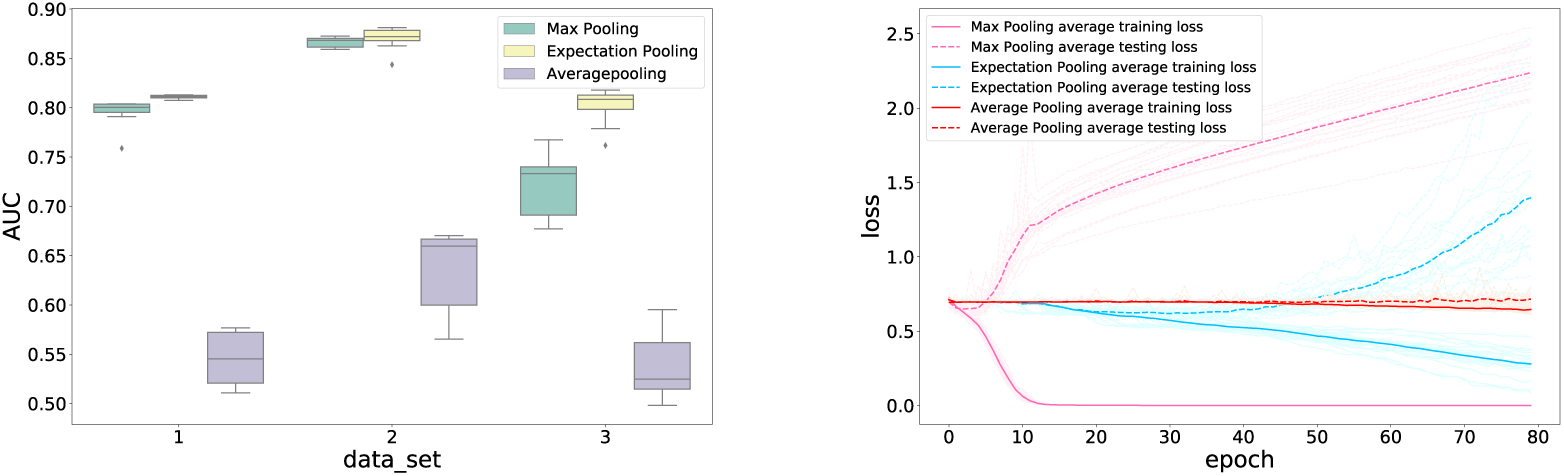
Expectation pooling performs better and is more robust to random seeds than are global max and average pooling (a), and expectation pooling suffers less from overfitting than global max pooling (b). Here (a) shows the AUCs of models with different pooling methods on the simulated datasets 1 (short motif), 2 (long motif) and 3 (mixed motifs). The mean AUCs on these datasets are 0.795774, 0.866507, and 0.720751 for models with global max pooling, 0.81092, 0.870577, and 0.801181 for expectation pooling, and 0.545254, 0.636738, and 0.53726 for global average pooling, respectively. (b) shows the learning curves for models with different pooling methods. The difference between training and testing loss for the model with expectation pooling is still moderate after 20–40 epochs; in contrast, the difference for the model with max pooling becomes very large immediately after 5 epochs. In addition, the model with global average pooling is difficult to fit and leads to low performance, while the methods with expectation pooling require more than 20 epoch to fit, and a slight overfitting occurs after 40 epochs. (a) The performances of models with different types of pooling on the (b) The training loss with different types of pooling on the simulated data simulated data

The performance improvement was especially evident on **simulated dataset 3** with a hard model, reflecting the superiority of expectation pooling in cases with complex motif settings. Clearly, on complex datasets, the huge fluctuation in the original model, it does not satisfy the need for fitting for low accuracy and it lacks robustness to initialization. On these simple simulated datasets, we can see that even a minor accuracy increase compared to the original model yields good performance.

### 3.2 Performance of real datasets

Having demonstrated the superior performance of expectation pooling on the simulated datasets, we next tested whether it could maintain this performance level on real-world cases. The neural network models differed only in their use of different pooling methods (i.e., global max pooling (baseline) and expectation pooling, respectively). We used the same model structures and parameter settings as in the preceding simulated unless explicitly stated otherwise. The window size of local max pooling was set to 10. The number of kernels varied from 8 to 128. The two models can be compared because expectation pooling does not increase the number of parameters.

The results show that when the number of kernels is limited (e.g., 8), the model with expectation pooling achieves a statistically significant improvement in AUC (one-sided Wilcoxon signed-rank test, p=4.01*e*^−84^); in particular, it achieved a better performance on 583(84.5%) of the datasets (see Table 2). However, its accuracy was lower on (107 approximately 15.5%) of the datasets, which does not match the theoretical analysis above. Because neural networks are not convex models, they do not necessarily obtain a global optimum. We selected the datasets on which our model’s performance was lower and chose several different random seeds for initialization. Subsequently, we found that the mean performance between the two models was almost identical (see the Supplementary Notes). In the DeepBind model, kernel length has a significant performance effects; thus, we considered another smaller representative number. We found that when the kernel length was set to 15, the baseline attained a better performance [Table 3], and our method achieved a greater improvement in average AUCs.

**Table 2.** Performances on real data

| Kernel number | Local window size | Maxpooling AUC(%) | ExpectationPooling AUC(%) | The percentage of improved dataset(%) | p value(wilcoxon) |
| --- | --- | --- | --- | --- | --- |
| 8 | 10 | 82.3 | 83.9 | 84.5 | 4.01E-84 |
| 16 | 10 | 84.1 | 85.5 | 88.6 | 4.853E-95 |
| 32 | 10 | 85.2 | 86.2 | 79.9 | 6.960E-72 |
| 64 | 10 | 86.1 | 86.5 | 65.3 | 5.10E-30 |
| 128 | 10 | 86.4 | 86.8 | 71.0 | 5.12E-33 |

**Table 3.** Performances on real data (kernel length=15)

| Option structure | Kernel number | Local window size | Maxpooling AUC(%) | ExpectationPooling AUC(%) | The percentage of improved dataset(%) | p value |
| --- | --- | --- | --- | --- | --- | --- |
| optional structure | 32 | 10 | 0.86174 | 0.87368 | 94.6 | 5.29E-103 |
| no optional structure | 32 | 10 |  | 0.87289 | 93.5 | 6.58E-98 |
| optional structure | 64 | 10 | 0.87094 | 0.88043 | 93.3 | 3.22E-96 |
| no optional structure | 64 | 10 |  | 0.87725 | 82.9 | 1.70E-74 |
| optional structure | 128 | 10 | 0.87579 | 0.88256223 | 87.4 | 1.02E-78 |
| no optional structure | 128 | 10 |  | 0.87832 | 71.1 | 4.37E-28 |

### 3.3 Varying the hyperparameters in our model

Next, we studied the effects of the hyperparameters on the performance of our model. In this section, we discuss the two newly added hyperparameters in our model.

#### 3.3.1 Varying *m*

The hyperparameter *m* controls the weights of each score: the larger *m* is, the greater the weights assigned to the high scores are during expectation pooling.

In this experiment, we also utilized one of the simulated datasets to determine the general rules of *m* for the models. From Figure 3, it is evident that after *m* is sufficiently large, the AUC of the model will decrease if *m* grows larger, which proves that the model with global max pooling is not optimal for motif finding (i.e., our model degrades to the baseline as *m* approaches infinity). In addition, a steep fluctuation is apparent when *m* becomes relatively small (i.e., between 1 and 5). We also note that, given that *m* is derived in the loss function, it could be learned from the data directly via the classical backpropagation algorithm, which would change *m* from a hyperparameter to a learnable parameter.

**Fig. 3.**
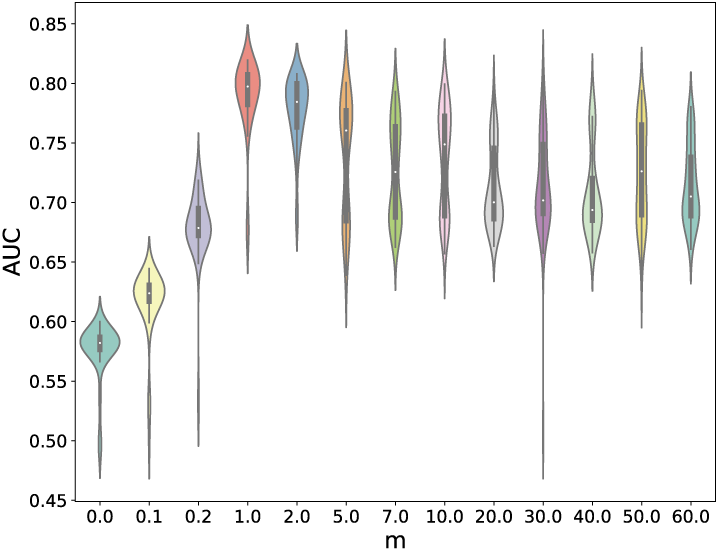
The AUCs of the model with varying *m* (violin plot). Note that when *m* = 0, expectation pooling is equivalent to average pooling, so the performance is worse than when *m >* 0. As *m* increases, the average AUCs also increase. When *m* reaches approximately 1, the performance is both stable and good. When *m* becomes too large, there is no difference between global max pooling and expectation pooling. Consequently, the performance degrades.

#### 3.3.2 Varying the window sizes of local max pooling

The window sizes used for local max pooling are also significant in our model. Because two regions of the motif cannot be located in too-close proximity, we need to calculate the max score of a local window, which means we must select a score to represent the whole window. In addition, a large window size requires fewer calculations. The other parameters (including batch size, kernel length, kernel number, batch size) are fixed to the same values shown in Table 1.

We trained models with different window sizes. the results in Figure 4 show that the AUCs generally increase when the window size increases from 1 to 15; subsequently, the model performance remains relatively stable. When the local window size is sufficiently large, expectation pooling degenerates into global max pooling. Consequently, we set the local window size to approximately 10 (this is approximately the median length of the motifs in the TRANSFAC database) or set it through validation.

**Fig. 4.**
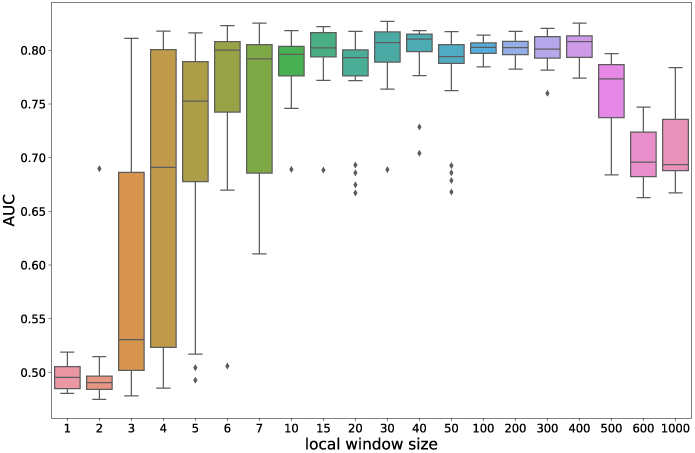
The AUCs of models with different local window sizes (box plot): The performance increases initially, but when the local window size exceeds approximately 5, the performance becomes almost stable. Finally, when the local window size approaches the sequence length (which means expectation pooling becomes equivalent to global max pooling) the performance equally as bad as that of global max pooling.

### 3.4 Model visualization

In this section, we show that the model with expectation pooling can recover the underlying motifs more accurately. Here, we used simulated data because we do not know the true motifs in real world datasets. We generated the sequence logos from kernels as described in Section 10.2 of the DeepBind Supplementary Notes(Alipanahi *et al.*, 2015). The best-recovered motifs (in the sense of information content) are compared to the true motifs utilized in the simulated data by calculating their similarity (E-value) with the Tomtom(Gupta *et al.*, 2007) algorithm.

The motifs recovered by our model and the model with global max pooling are both aligned to the true motifs. (Figure 5). However, based on the E-value, we found that the sequence logos generated by our model are more informative and better match the ground truth. This result demonstrates that our model is able to find more accurate motifs. In addition, expectation pooling can clearly distinguish the motif regions from other regions (obeying the background distribution). In Figure 5 (b), the motif recovered by the origin model contains obvious noise in addition to the eight positions corresponding to the true motif, while in Figure 5(d), the noise is not obvious in our model.

**Fig. 5.**
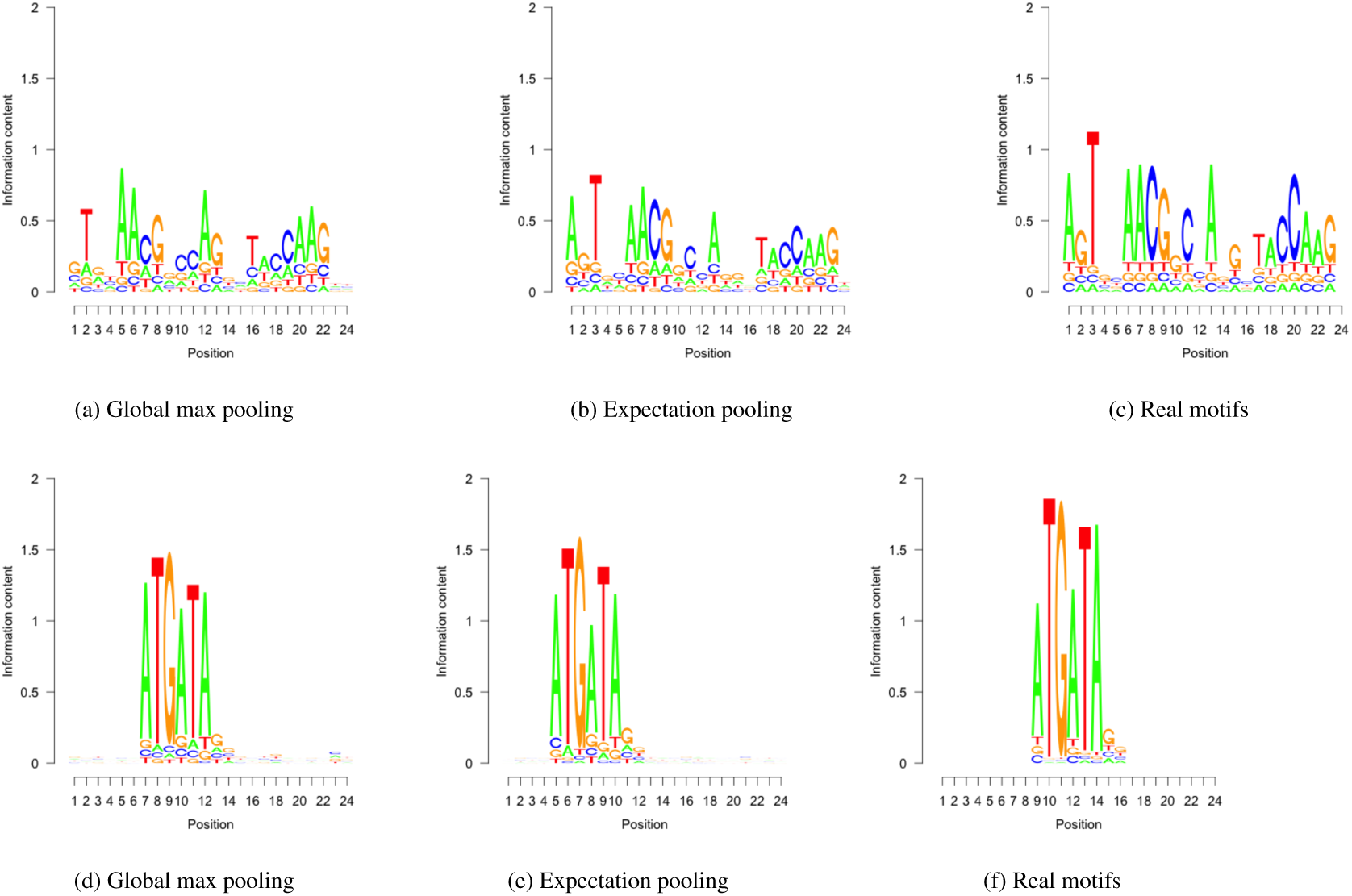
Motifs recovered by our model (middle row) and by CNNs with global max pooling (top row) compared to the true motifs (bottom row). As a result, the E-values of the motifs recovered by our model are 2.04 × 10^−^19, 1.66 × 10^− 8^, respectively, while the ones recovered by CNNs with global max pooling are 1.59× 10^−17^, 2.55 × 10^−6^, respectively

## 4 Discussion

### 4.1 Expectation pooling as the E-step in an (object-optimized) EM algorithm

In the context of sequence motif detection (see Supplementary Notes for a brief summary on the typical CNN architecture used for motif detection), expectation pooling can be interpreted as an (object-optimized) EM algorithm.

Briefly, given a particular motif represented as as a position weighted matrix(Stormo, 2000) (PWM) M, the i-th sequence *X*_*i*_ (positive sample) and the motif location *Z*_*i*_ = *j*(*Z*_*i*_ = *j* if motif starts at position *j* in sequence *i*), we obtain

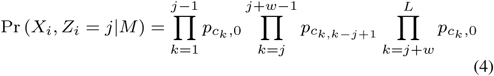

where *c*_*k*_ is the character at position *k* in sequence *i*, 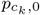 is the probability of *c*_*k*_ in the background distribution, 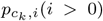 is the probability of *c*_*k*_ in the distribution of the i-th position in the motif region. Here, if PWM *M* is given, we have the following E-step(Buhler and Tompa, 2002; Dempster *et al.*, 1977; Lawrence and Reilly, 1990):

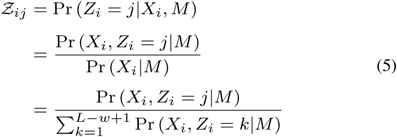

in which *Z*_*i*_ is the latent variable (i.e., the start position *Z*_*i*_ = *j* is unknown) and Ƶ _*ij*_ represents the likelihood of *X*_*i*_’s motif starting at position *j*. The Ƶ _*i*_ = {*Ƶ* _*ij*_} represents the distribution of starting position of *X*_*i*_’s motif. Next, we update PWM using the M-step and iterate until convergence.

Given the exact transformation between the convolutional layer kernels and PWM(Ding *et al.*, 2018), the log-likelihood of the resulting PWM of any DNA sequence is exactly the sum of a constant and the convolution of the original kernel on the same sequence.

The formula is as follows:

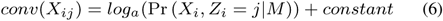

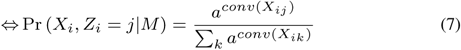

where *conv*(*X*_*ij*_) represents the convolution result of the subsequence starting at position *j* of sequence *X*_*i*_ with some kernel and is the same as *A*_*i*_, the input to the pooling layer, and *a* is a given constant.

As a result, the pooling layer input is a score vector equivalent to the log-likelihood, namely, the larger the score, the more “similar” the motif is to the specific sequence fragment it aligns to and the more likely it is to have a positive label.

Next, we derive that our expectation pooling is equivalent to the expectation of the log probability of the motif in sequence *X*_*i*_ given *a* = *e*^*m*^:

#### pooling value

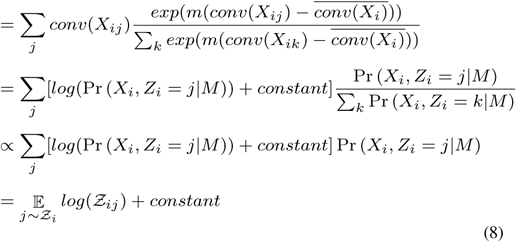

Eq.8 shows that the output of the expectation pooling is proportional to the expectation of the motif in a sample when *m* is appropriate from the statistical model perspective. Similar to the E-step of the EM algorithm, which utilizes a distribution of Ƶ_*i*_ and the calculation expectations combined with the distribution, expectation pooling not only considers the highest peak of the score curve but also considers every other score (see Figure 1(b)) but with less emphasis. In contrast, max pooling considers only the highest score, which can be affected by false positives caused by random fluctuations of the background distribution (see Figure 1(b)). Given the short lengths of sequence motifs, there is a high probability that a high score will correspond to the real PWM by coincidence given the background distribution for negative samples.

However, calculating the expected value directly leads to underestimation and requires excessive computation, which is not desirable if many more kernels are utilized. To solve this problem, we conduct local max pooling (i.e., the first sublayer of expectation pooling) before calculating the expected value. Furthermore, local max pooling filters the majority of small scores, which offsets the disadvantage of taking the expectation of all the scores and leads to a phenomenon referred to as “trimming the hills and filling the valleys.”

After local max pooling, the probability (i.e., the weight) set for each score is an exponent (i.e., weight, 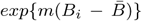 without normalization, in which *m* is an adjustable parameter. Thus, if *m* tends toward infinity or zero, expectation pooling will obviously be reduced to either max pooling or average pooling, respectively, which also shows that our expectation pooling is an improvement from a probability viewpoint, but differs from stochastic pooling due to its forms of expectation.

### 4.2 Benefits of expectation pooling

We know that the Gaussian mixture model is often referred to as a soft clustering method, while K-means is comparatively hard (Friedman *et al.*, 2001; MacQueen *et al.*, 1967) because the Gaussian mixture model does not directly yield sample labels according to the minimal distance; instead, it applies labels from the viewpoints of probability and expectation. Actually, expectation pooling inherits ideas from the Gaussian mixture model and it yields a “soft” maximum of the scores while pooling.

Expectation pooling provides two main benefits. On one hand, expectation pooling considers all the scores rather than only the maximal score, which improves model robustness. This point is significant because a motif-detection model is a probabilistic model with high randomness. For example, for one positive and one negative sample, if the highest scores of a kernel of two models are nearly the same, the outputs of expectation pooling are obviously distinguishable when the positive sample has more high scores. On the other hand, expectation pooling can be shown to play a role in reducing overfitting. Overfitting is a modeling error that occurs when a function is too closely fit to a limited set of data points. It has been shown that max pooling has the drawback of overfitting easily, while average pooling avoids this problem. Consequently, our neural network models do not overfit easily without a dropout layer(Srivastava *et al.*, 2014) for regularization, which is essential for DeepBind. Actually, our method of pooling combines average pooling with max pooling in a reasonable way whose probability interpretation corresponds to that of the EM algorithm.

The first sublayer of expectation pooling is also essential not only for reducing calculation time but also for overcoming the drawback of the underestimation of the second sublayer, which can occur because many zero elements are included in the scores after ReLU activation. From a statistical viewpoint, it is abnormal to regard two close positions as two different motif start positions and to consider their scores twice. As a result, expectation pooling is a balanced global max pooling with clear interpretability.

### 4.3 The effect of kernel numbers on motif inference

By experimenting with different hyperparameters, Zeng *et al.* (2016) demonstrated that including more kernels in convolutional layers can lead to better performance. However, although many kernels may be utilized, the truly effective and significant kernels are limited after training according to model visualization. Considering the limited number of effective kernels, using fewer kernels is reasonable from a model perspective, considering the number of required calculations and model visualization. However, extra kernels may affect the optimization process by avoiding premature convergence due to becoming trapped in a local minimum of the loss function. The results show that increasing in the number of convolutional kernels has a smaller impact on performance, but is significant in the model with global max pooling. Therefore, our model with expectation pooling is robust to the number of kernels (also see the Supplementary Notes).

### 4.4 Optional structures

We also notice that the expression in the second sublayer (optional structure) can be modified as follows:

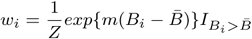

where 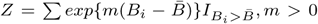 and *I* represents an indicator function to increase the sensitivity for motifs with only a few conserved residents (i.e., when the pooling layer input consists of a few large numbers but many small numbers. In Figure 6, expectation pooling attains a better performance on 94.6% of the datasets when the optional structure contains 32 kernels; these configurations resulted in the best AUCs among the different hyperparameter values[Table 3].

**Fig. 6.**
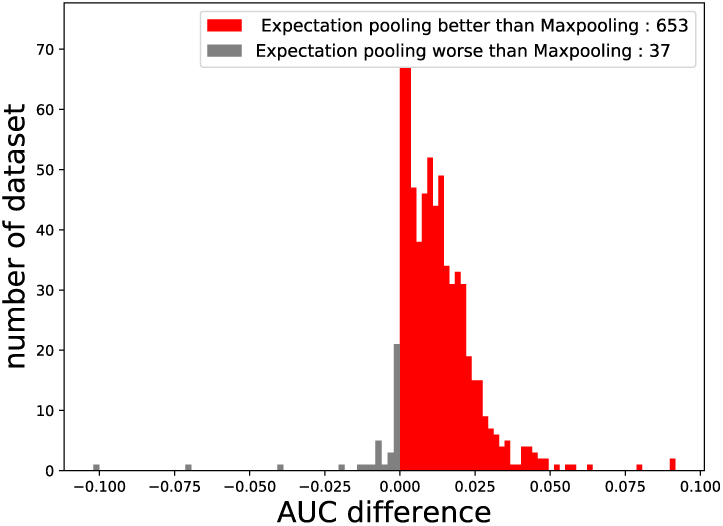
The performance of our model utilizing expectation pooling on real datasets, where the window size = 10, m = 1, kernel number = 8, kernel size = 15 and the optional structures are utilized. Expectation pooling increases the AUC on real datasets. The x axis shows the AUC difference under expectation pooling and max pooling. The models with expectation pooling are better than are the models with max pooling on 653 datasets but worse than the models with max pooling on 37 datasets. This figure clearly shows that expectation pooling achieves better performances

Moreover, a hidden layer with a dropout layer can be introduced after expectation pooling to increase model learning ability. A dropout layer is applied to a hidden layer to randomly mask portions of its output to avoid overfitting. The introduction of the hidden layer sacrifices some portion of model interpretability but helps in modeling complex datasets without prior model assumptions.

### 4.5 Summary

In this paper, we introduced a novel pooling method, termed expectation pooling, to improve the performance of DNA sequence-related prediction tasks. Expectation pooling is divided into two sublayers, a local max pooling layer and a “dense” layer without additional hyperparameters (other than *m*). Expectation pooling is a novel combination of average pooling and max pooling, and its performance in other fields should be investigated. In this paper, we show that expectation pooling improves model performance compared with global max pooling. Our method improves the performance only from the aspect of the meaning of the model results—without increasing parameters or requiring data augmentation—and allows the neural network model to be related to a probabilistic model.

In addition to the experimental results presented here, expectation pooling is more suitable for the model of motif finding from two perspectives: those of deep learning and those of statistical models. From a statistical perspective, expectation pooling actually calculates the expectation of kernel scores; it is evident that expectation pooling matches what MEME does better than does max pooling. From the of deep learning perspective, the drawback of max pooling is overfitting and overestimation, while the drawback of global average pooling is underestimation. Thus, we combine these two pooling methods to introduce our new expectation pooling method. Additionally, considering all the scores improves our model’s robustness. From the analysis above, the experimental performances are expected and can be interpreted from many aspects.

We also considered probabilistic pooling methods, which yield a random largest score rather than a weighted average of the large scores. Probabilistic pooling means that we consider the weight to be the probability selected as the final output. Actually, our expectation pooling uses the output expectation for convenience and robustness without randomness, which is how ‘expectation’ is formed. Thus, our results tend to be more stable and robust. However, we believe that the probabilistic pooling mentioned above can be utilized during the training process, for example, in dropout(Srivastava *et al.*, 2014), to enhance regularization and prevent CNNs from overfitting.

The motif finding problem remains unsolved. Deep learning is magical when dealing with large datasets and intricate structures, and it has dramatically improved the state-of-the-art in many fields. Neural networks have achieved numerous successes, such as DeepBind(Alipanahi *et al.*, 2015) for motif inference. Nevertheless, despite its great achievements, deep learning is also blamed for its lack of interpretability(Zou *et al.*, 2018; Castelvecchi, 2016). Recently, many novel neural networks(He *et al.*, 2016) have been proposed based on intuition rather than through logical derivation. Because the original models we utilized are simple and related to the probabilistic model of motif finding, expectation pooling is a natural improvement informed by the probabilistic model utilized in MEME(Bailey *et al.*, 2006). Obviously, global max pooling does not match the statistical model particularly well; thus, we add the idea of expectation to the model. As mentioned before, expectation pooling is a promising modification of global max pooling that transitions from “hard” pooling to “soft” pooling from a statistical viewpoint. We believe that the problem of motif finding implies a simple statistical model and consequently, that simple neural network models can be applied to analyze this problem. Under simple models, statistical methods can be utilized reasonably while still obtaining some seemingly magical performances. Finally, the interpretation of expectation pooling enhances model understanding; therefore, we recover more accurate motifs as expected.

Recently, many works have been conducted to investigate the interpretation of neural networks and to improve the prediction accuracy in the motif-finding field(Cao and Zhang, 2018; Pan and Shen, 2018; Pan *et al.*, 2018; Zuallaert *et al.*, 2018). These works have given us more knowledge about the CNNs utilized in these models. Moreover, many recent statistical methods such as clustering have also been used to construct successful motif-finding applications(Munteanu *et al.*, 2018). These works inspired us to combine statistical models with motif finding rather than experiment blindly with new deep learning models. From a statistical viewpoint, many novel and reasonable technologies may be proposed for this problem and utilized in deep learning, such as dropout and expectation pooling. Furthermore, our pooling method has a significant statistical relationship with the EM algorithm, which gives our model better interpretability. Better interpretability, we believe is more impactful than experimental performance because it allows us to understand biological models from their statistical models and provides interesting ideas from a probability viewpoint. Our work in this paper is generally instructive for applications of statistical methods in the motif-finding field. We believe that statistical methods combined with deep learning forms a substantial advancement by will making deep learning an even more powerful tool for bioinformatics.

