## Supplementary material for "Expectation pooling: An effective and interpretable pooling method for predicting DNA-protein binding"

### Supplementary Notes

#### 1 Baseline

Each DNA sequence is transformed into one-hot format. Specifically, it's turned into $4 \times L$ matrix where each base pair in the sequence is denoted as one of four one-hot vectors $[1, 0, 0, 0]$, $[0, 1, 0, 0]$, $[0, 0, 1, 0]$ and $[0, 0, 0, 1]$. The first layer is a 1-D convolutional layer with ReLU activation. The second layer is a global max pooling layer (i.e., each kernel yields one value). The last layer is a fully connected layer with one output. The last layer is a fully connected layer with one output.

#### 2 Supplementary Figures

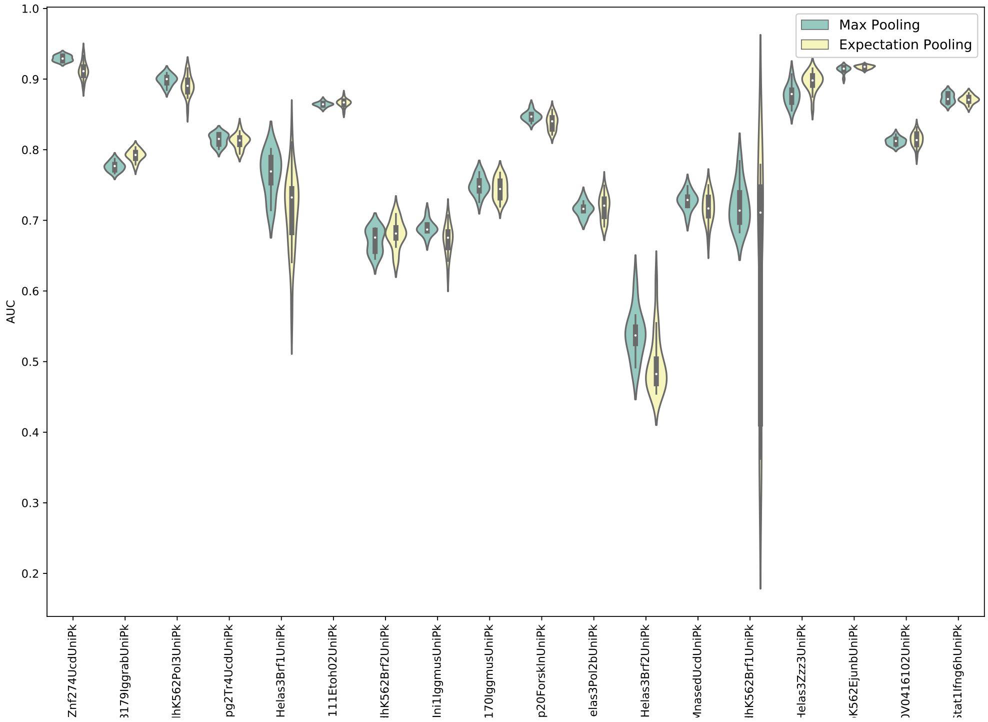


Figure S1: We selected the datasets for which expectation pooling was below 0.01 and executed them 10 times with different random seeds to determine whether the poor performance was consistent. As shown, on these datasets, expectation pooling is not always worse than the max pooling; in fact, the mean performance of the two models is almost identical.

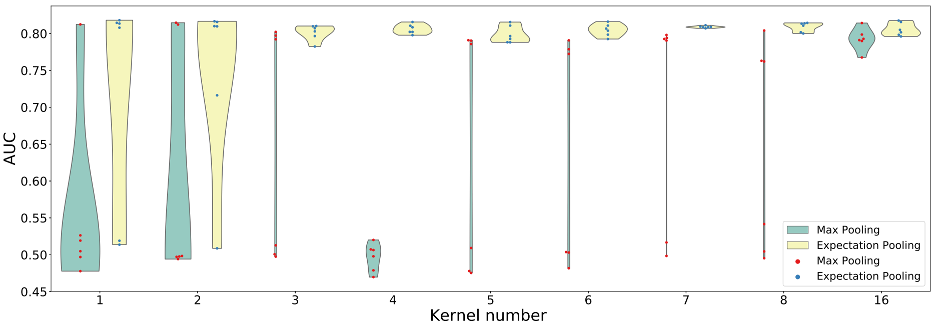


(a) Simulated data 1

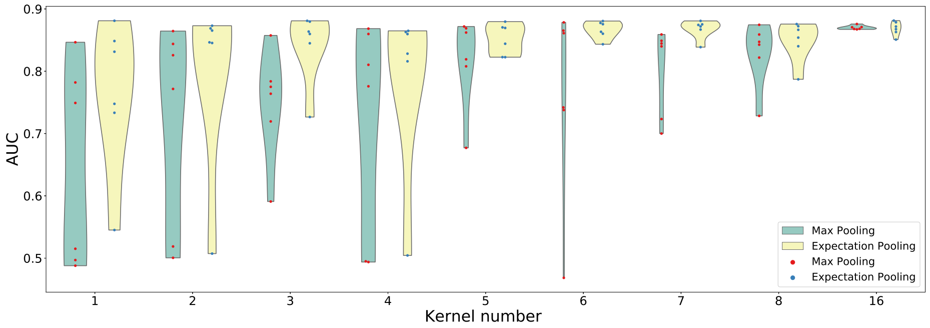


(b) Simulated data 2

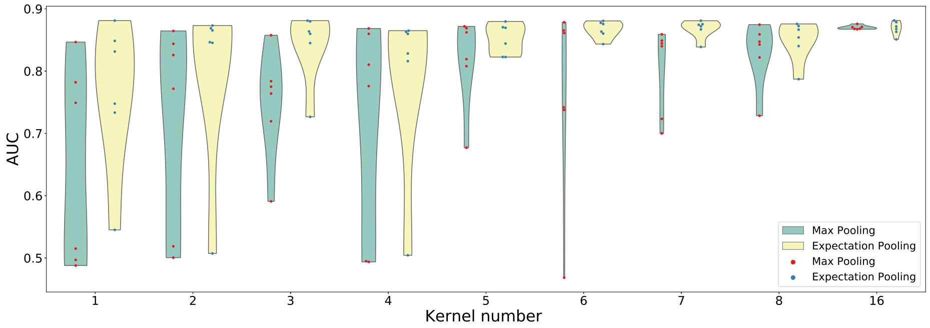


(c) Simulated data 3

Figure S2: The AUCs of models with different kernel numbers on all the simulated data sets. With limited kernel numbers, global max pooling tends to obtain more robust and better results.
